# Dynamic connectivities of plant metacommunities at a millennial time-scale: the Beringia testbed

**DOI:** 10.1101/2025.08.19.671027

**Authors:** Ulrike Herzschuh, Kathleen R. Stoof-Leichsenring, Luca Zsofia Farkas, Ying Liu, Sisi Liu, Thomas Böhmer, Izabella Baisheva, Boris K. Biskaborn, Jérémy Courtin, Bernhard Diekmann, Laura S. Epp, Barbara von Hippel, Sichao Huang, Yongson Huang, Weihan Jia, Darrell S. Kaufman, Hanno Meyer, Martin Melles, Stefano Meucci, Luidmila A. Pestryakova, Luise Schulte, Bernd Wagner, Zhigang Wang, Evgenii S. Zakharov

## Abstract

Ecological connectivity shapes ecosystem responses to climate change and is thought to underpin stability, yet its millennial-scale dynamics remain poorly resolved. We asked how spatial and temporal connectivity of plant metacommunities changed over the last 40 ka and which processes drove it. We analysed and compiled plant sedimentary DNA from 20 lake cores across Beringia (Siberia, Alaska) to investigate community dynamics and, for a high-resolution subset, applied beta- and zeta-diversity to track connectivity. Vegetation changed coherently across the glacial–Holocene transition, with trait shifts mirroring functional composition. Connectivity peaked during the late MIS3 and Last Glacial Maximum—likely aided by the Bering Land Bridge, mass effects and facilitation—collapsed during the Deglacial with rapid turnover, and rebounded in the Holocene as shrub and boreal communities expanded. Temporal zeta within sites exceeded spatial zeta, indicating strong local persistence and resilience. Tundra sites uninvaded by forest maintained continuous species pools. Overall, these patterns underscore the value of a metacommunity perspective for assessing millennial-scale connectivity changes.

## Introduction

Ecological connectivity fundamentally shapes how ecosystems respond to climate change through shifts in local community composition (*1*). In this context, resilience is often enhanced by spatial connectivity among habitats, which facilitates species migration (*2*), and by temporal connectivity (continuity), which supports species persistence (*3*). However, spatially or temporally isolated communities can also exhibit resilience, as reduced connectivity limits colonization by new species, potentially preserving established community structures (*4*). Thus, despite recognizing the general importance of connectivity, the precise roles of spatial and temporal connectivity in shaping community responses to climatic fluctuations remain poorly explored by empirical studies.

The metacommunity concept provides a valuable framework for investigating the role of connectivity in shaping species assembly processes under climate change (*5*). It describes communities as networks of local habitats interconnected through species dispersal. Within this framework, species interactions and species-specific responses to local environmental conditions—largely determined by species traits and environmental niches—play key roles in shaping community composition and dynamics. However, assessing the metacommunity concept’s applicability to species assembly requires empirical evaluation under extensive natural spatiotemporal environmental variability (*6*).

Beringia, encompassing eastern Siberia and Alaska, represents an ideal testbed for exploring ecological concepts about long-term metacommunity connectivity, due to its unique geographic setting (*7*) and heightened sensitivity to climate fluctuations (*8*, *9*). Although paleovegetation studies have often targeted compositional shifts and community reorganizations (*10*, *11*), changes of the ecological connectivity between eastern and western Beringia in relation to the formation during the late Marine Isotope Stage (MIS) 3 and subsequent flooding of the Bering Land Bridge at the beginning of the Holocene have been less well investigated using paleodata (*12*). So, it remains largely unknown how changing connectivity impacted the composition of the glacial steppe biome (*13*). Furthermore, it is uncertain which roles did species traits—such as dispersal capacity and life-history strategies—play in shaping communities under strong postglacial climate change, particularly in relation to deterministic versus stochastic processes. Plant-mammal interactions changed (*14*) with the Deglacial megaherbivore loss, as did plant-plant interactions with woody taxa expansion during the Holocene (*15*). However, how these factors of species-assembly influenced ecological connectivity both spatially and temporally across Beringia, remains unclear. Resolving these uncertainties necessitates robust paleoecological data and advanced analytical methods capable of capturing intricate spatio-temporal ecological dynamics.

High-resolution lake sediments offer robust environmental archives for reconstructing vegetation dynamics extending beyond the Last Glacial Maximum (*16*). Metabarcoding of sedimentary DNA using the plant gh marker captures community composition at a higher taxonomic resolution and more local area compared to pollen analysis (*17*, *18*), which is traditionally used for past vegetation assessment. Plant metabarcoding studies have already provided critical insights into postglacial species assembly in northern Europe (*19*), shifts in species richness relative to mean range size (*15*), temporal patterns of potential plant extinctions (*20*), and Beringian vegetation change (*21*, *22*), among others. Diversity measures are particularly informative within metacommunity contexts (*23*). Alpha diversity reflects local community diversity such as richness at a given point in time. Beta diversity quantifies compositional turnover between communities, for example capturing how species similarity declines with increasing geographic distance—a phenomenon known as distance decay (*24*)—highlighting spatial connectivity or isolation among communities. Zeta diversity (*25*), measures species retention across multiple sites at a specific time or at one site through multiple time periods, thereby allowing direct comparisons between spatial and temporal connectivity in species pools. Applying these measures within the context of past climate change enables the examination of site-specific and regional species pools and their interconnectedness. This approach provides a robust framework for investigating essential ecological processes such as dispersal, colonization, extirpation, and community turnover. Integrating trait-based data, including growth form, dispersal capacity, and the competitor-ruderal-stress-tolerance strategy (CSR) (*26*), further enables deeper insights into the underlying mechanisms driving species assembly.

This study employs plant metabarcoding of sedimentary ancient DNA from multiple lake sediment cores across Beringia to investigate plant community dynamics focusing on the last 28,000 years (28 ka). By applying advanced diversity metrics, we explore how climatic change has shaped the connectivity of plant metacommunities in space and time. Our findings indicate that Beringian vegetation responded coherently to the Pleistocene–Holocene transition, highlighting that substantial compositional shifts are also reflected in trait shifts. Enhanced connectivity across Beringia during the Last Glacial Maximum (LGM), likely driven by the establishment of the Bering Land Bridge and supported by mass effects and facilitative plant interactions, contrasts sharply with the rapid community turnover and disrupted connectivity observed during the Deglacial period. The Holocene period subsequently saw increased regional connectivity, marked by stable community structures dominated by boreal forests and shrubs. Additionally, our analysis reveals long-term resilience in tundra sites not invaded by forest, emphasizing the critical role of temporal connectivity in supporting taxa persistence along with shaping plant community dynamics under changing climates.

## Results

### SedaDNA metabarcoding data set and spatio-temporal pattern of plant communities

Plant metabarcoding analyses of 1017 lake sediment samples originating from 20 sediment records (Fig. 1a and table S1) yielded, after filtering, a total of 226,261,847 reads which could be assigned to plant taxa from the regional SibAla_2023 reference database with 100%-identity. The dataset includes 482 amplicon sequence variants (ASVs); of which 46% are assigned to species level and 38% to genus level.

**Fig. 1.**
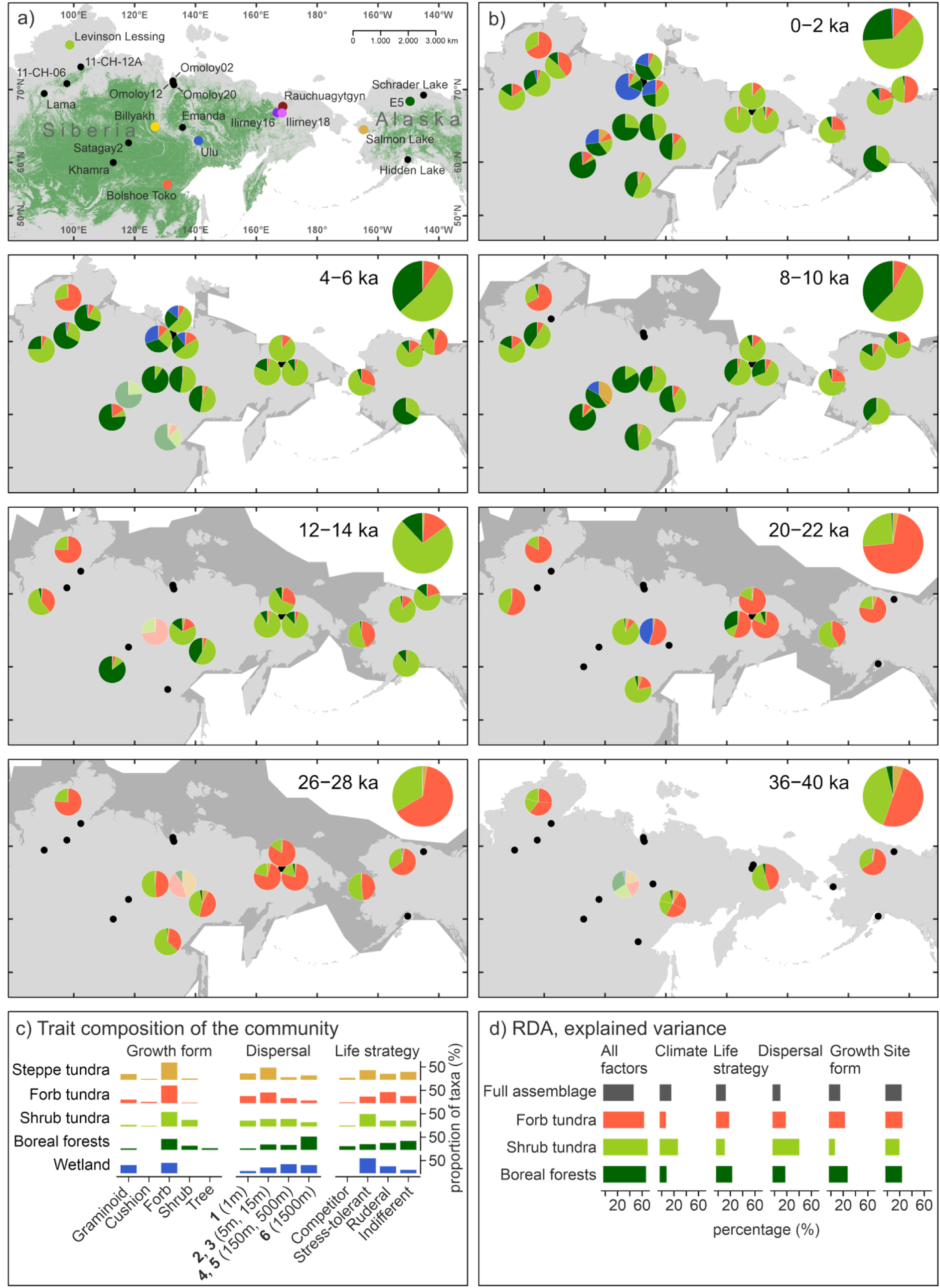
Site locations, community composition and trait information. (**a**) Location of the study sites investigated for sedaDNA plant metabarcoding, all of which were used to examine spatio-temporal changes in meta-community composition. From these 20 sites, nine long-term and high-resolution records (indicated by large-sized colored dots) were selected to study detailed diversity changes over the past 28 ka. The forest cover in panel a was derived from the ESA CCI Landcover Classes for needleleaved forest (*105*). (**b**) Portion of metacommunities per site for selected time-slices over the past 40 ka. Big pie-charts in the upper right corner represent median-composition across all sites. Transparent piecharts with potentially poor data quality as traced by unusual low richness. The dark gray shaded areas represent unflooded landmass for the respective time-slices which are flooded at present (see details about data source in method section) (**c**) Trait composition for five metacommunities: growth form, dispersal classes with 99 percentile dispersal distances (*66*), and life strategy (*26*). (**d**) Explained variance by changes of single traits and site assessed for full taxa assemblage and major metacommunities from a series of redundancy analyses (RDA).

Community identification using co-occurrence network analyses of plant assemblages in the original samples yielded five communities composed of 395 of the 482 taxa. The five communities are steppe tundra, forb tundra, shrub tundra, boreal forest, and wetland communities (see taxa composition in Fig. 1c, fig. S1a, b, and table S2). The community composition for each record for selected time-slices is shown in Fig. 1b and fig. S2a.

The steppe tundra community comprises 80 taxa, consisting mainly of forbs and a relatively high number of graminoids compared to other communities (Fig. 1c and table S2). Overall stress-tolerant and locally dispersing taxa dominate. Pooideae*, Potentilla*, Apioidae, *Koenigia*, *Sorbaria sorbifolia,* and *Pulsatilla dahurica* are characteristic taxa. The community generally covers a low proportion of reads, but is more common during the glacial period, particularly in Yakutia (Fig. 1b).

The forb tundra community includes 101 taxa, predominantly of forbs (Fig. 1c and table S2). Most characteristic are *Papaver*, *Bistorta vivipara*, *Oxyria digyna,* and *Myosotis alpestris*. The community also includes many cushion plants such as *Eritrichium* and *Saxifraga* and dwarf shrubs such as *Dryas*. It is the only community with a substantial proportion of bryophyte taxa (fig. S1b). Most taxa are characterized by short dispersal distances. The community has a high proportion of ruderal taxa (Fig. 1c). Overall, this is the most common community in our records (Fig. 2a). Before 14 ka BP, this community dominated most sites and the Lake Levinson Lessing record is characterized by this community at >60% throughout (Fig. 1b).

**Fig. 2.**
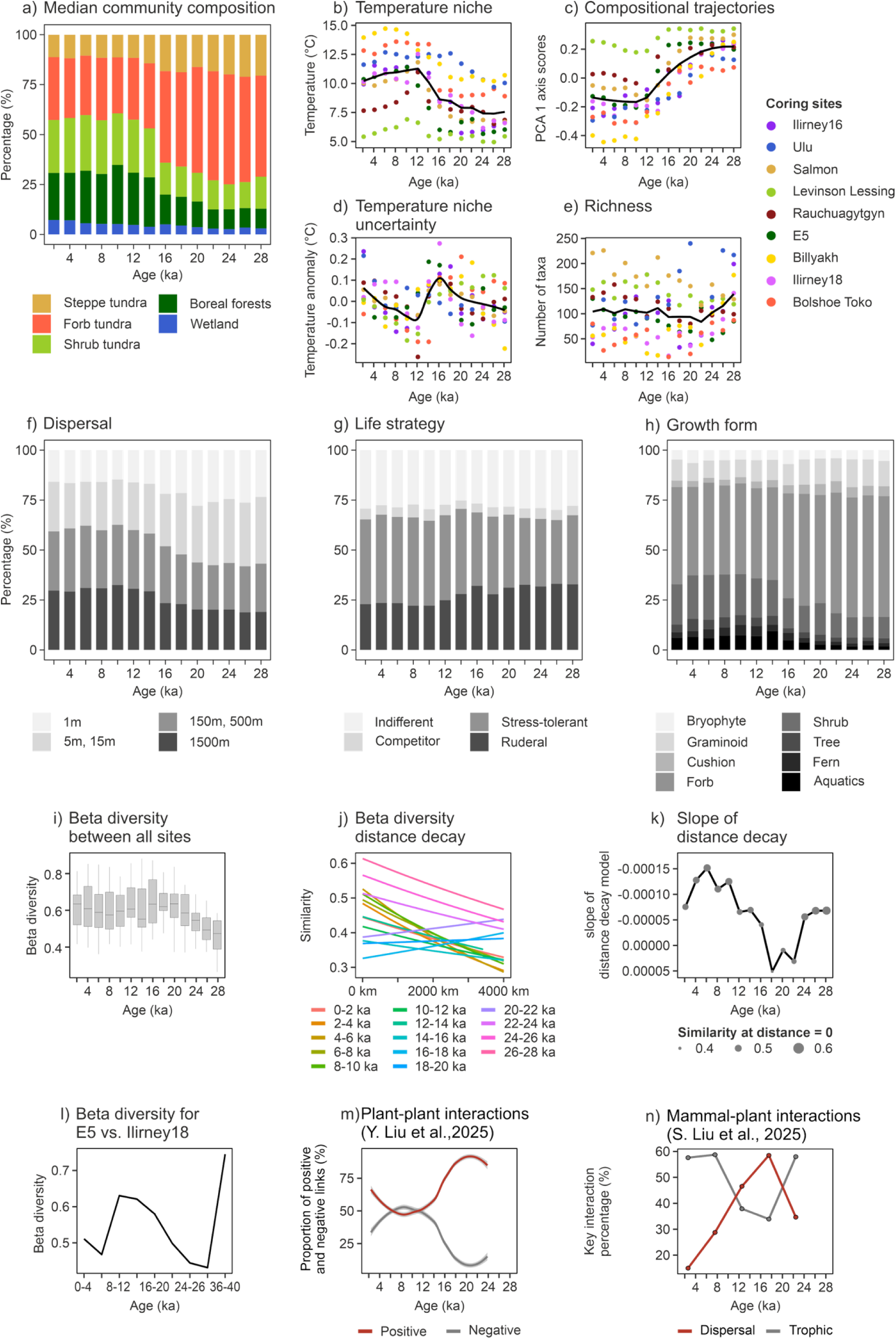
Trait trait, diversity time-series and ecological changes over the last 28 ka. (**a**) Metacommunity composition inferred from nine cores covering the past 28 ka. (**b**) and (**d**) Summer temperature niche and uncertainty reconstructed from sedaDNA. **(c)** PCA scores for the nine cores covering the past 28 ka. (**e**) Alpha diversity, representing site species richness per 2 ka time-slice for the nine cores covering the past 28 ka. (**f**–**h**) Trait composition for the nine cores covering the past 28 ka. (**i**) Beta diversity between the nine cores. (**j**) Beta diversity distance decay. (**k**) Slope of distance decay. The size of the points represents the first distance decay model parameter (i.e. the similarity at distance = 0) and the line indicates the second distance decay model parameter. (**l**) Beta diversity comparison between Lakes E5 and Ilirney18. (**m**) Proportion of positive and negative links in plant-plant interactions indicating that facilitation dominated during the Glacial and competition during the Holocene (*47*). (**n**) Proportion of mammal-plant dispersal and trophic interactions from network analyses of palaeogenomic data (*49*).

The shrub tundra community includes 76 taxa mainly characterized by dwarf shrubs like Saliceae, *Rhododendron tomentosum*, *Pyrola*, *Vaccinium vitis-idaea*, *Empetrum nigrum*, *Chamaedaphne calculata*, *Arctous rubra*, and *Cassiope tetragona* and several forbs. All dispersal modes are about equally represented and stress-tolerant taxa dominate (Fig. 1c). The share of shrub tundra before 16 ka is high at southern and central Yakutian sites (>50%) but low in the north (Fig. 1b). Most sites have high values during the Bølling–Allerød period (15–13 ka BP). While the shrub tundra proportion stays at high values of around 70% in the north it declines to less than 30% in the south at the transition to the Holocene at 11 ka BP.

The boreal forest community comprises 116 taxa among them all tree taxa including *Larix*, *Pinus*, and *Picea* and shrubs such as *Alnus alnobetula*, *Betula*, *Ribes*, *Sambucus,* and *Rubus*. Characteristic forbs are *Chamaenerion angustifolium* and Rosoideae. It is the community with the highest proportion of ferns (fig. S1b) including Lycopodioideae, *Thelypteris palustris,* and *Equisetum*. Compared to other communities, the boreal forest community has many taxa with long dispersal distances (Fig. 1c). Taxa with an indifferent CSR strategy dominate (Fig. 1c). Boreal forests make up a low proportion during the glacial period but increase to >30% in southwestern and central Yakutia (Billyakh and Bolshoe Toko, Fig. 1b). This community comprises most submerged aquatic taxa (fig. S1b) including *Potamogeton perfoliatus* and other *Potamogeton* taxa, *Ceratophyllum demersum*, *Callitriche hermaphroditica,* and *Myriophyllum,* which likewise markedly increased during the early Holocene.

The wetland community comprises 22 taxa including bryophytes (fig. S1b) (e.g. *Sphagnum* and *Aulacomnium*), graminoids (e.g. *Eriophorum* and *Carex aquatilis*) and forbs (e.g. *Tephroseris*). Most taxa are stress-tolerant and have a wide dispersal range (Fig. 1c). The wetland community comprises a rather low proportion of the vegetation and reaches >10% mainly after 6 ka BP in records from the north Siberian lowlands (Fig. 1b).

### General trends in community and trait compositions

Temporal changes of plant community characteristics are summarized in Fig. 2a–h, fig. S2. Typically, time-slices from the LGM and early Deglacial are dominated by forb tundra (Fig. 2a); have a high share of taxa with low temperature niches (Fig. 2b); many graminoid, cushion, and forb growth forms (Fig. 2h); and relatively many taxa with a ruderal life strategy (Fig. 2g) and non- or short-distance dispersal, where mode 1 and modes 2/3 dominate (Fig. 2f). The niche uncertainty is highest during the early Deglacial (Fig. 2d). Shrubs and stress-tolerant taxa increased in most records in timeslices from the Bølling–Allerød, while the Holocene is characterized by high percentages of trees, shrubs, and stress-tolerant taxa (Fig. 2g and h).

We performed a principal component analysis (PCA) on taxonomic composition from all records with samples aggregated into 2-ka time-slices. The taxa of the boreal forest and shrub tundra communities are located on the right side of the ordination biplot and the taxa of the tundra communities on the left side (fig. S3). The site-trajectories show coherent trends along the first PCA axis, explaining 18% of the dataset, with maximum values around the LGM and minimum values during the early Holocene (Fig. 2c). Increasing values of the second axis reflect a gradient from sites in Alaska/Chukotka to central Yakutia.

Redundancy analyses (RDAs) of those nine cores covering the last 28 ka yielded site as explaining the highest variance (Fig. 1d and table S3). Among the traits, climate niche best explains the compositional signal. Although ‘site’ explains a high share in all RDAs applied to the community level data, it varies with which trait explains the variation best (table S3). An RDA biplot is provided in fig. S4.

### Spatio-temporal patterns of alpha and beta diversity

Typically, taxa richness was highest during early LGM (∼150 taxa), lowest during the early Deglacial at ca 19 ka (∼100 taxa), and at intermediate levels since the Bølling–Allerød warming (Fig. 2e). However, trends in richness vary considerably at individual sites. (A plot with rarefied richness is provided in fig. S5 showing similar trends.)

To explore compositional dissimilarities between eastern and western Beringia over the past 40 ka, we performed beta diversity analyses specifically on sediment cores from Ilirney18 (EN18208, Chukotka, Siberia) and E5 (Alaska). These analyses show highest beta diversity during 40–36 ka BP, while lowest diversity occurred between approximately 32 and 24 ka BP (Fig. 2i).

Beta diversity assessments reveal low average beta diversity among sites and high compositional similarity at short spatial distances with only a weak distance-decay pattern during the 27–23 ka BP period. In contrast, between 21 and 17 ka BP, overall similarity is generally low (high beta diversity), showing no clear relationship with spatial distance (Fig. 2j). Intermediate beta diversity values characterize the period from 15 ka BP to the present, accompanied by a pronounced distance-decay relationship, most notably at around 5 ka BP.

### Zeta diversity pattern per time-slice and site

Zeta diversity—the number of taxa shared across a given number of sites or time-slices (zeta order)— along with richness-normalized zeta diversity and associated species retention curves (ratios between consecutive zeta orders), are presented in Fig. 3a–h. Zeta diversity indicates substantial taxon sharing among sites; for instance, Zeta₅, representing taxa shared across five of the nine coring sites within a given time-slice, ranges from 10 to 40 (Fig. 3a).

**Fig. 3.**
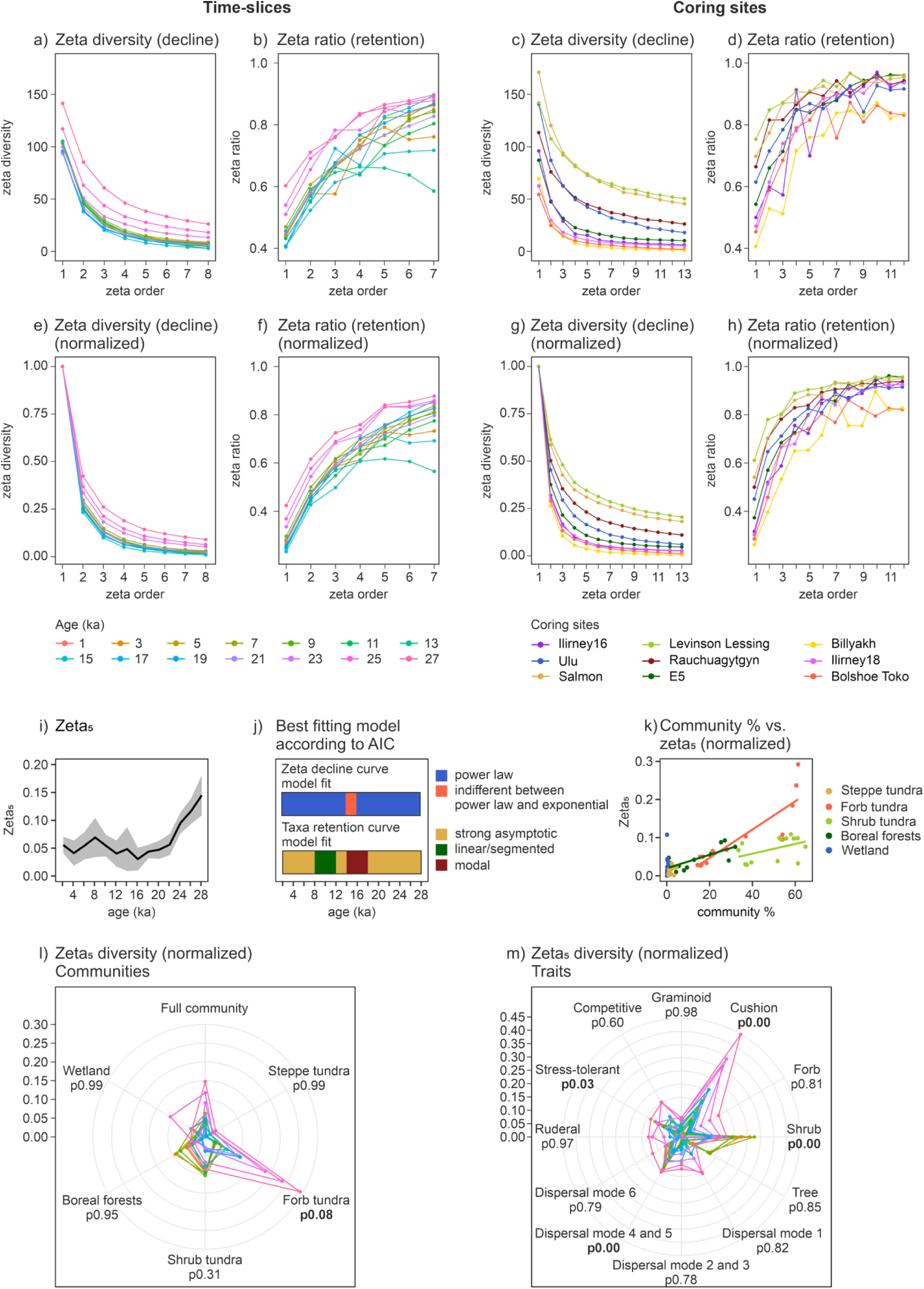
Zeta diversity across time-slices and coring sites of full community. **(a**–**j), with Zeta_5_ of communities and traits (k**–**m).** (**a–b**) Zeta diversity decline and ratio for time-slices and (**c–d**) for coring sites. (**e–h**) Normalized zeta diversity decline and zeta ratios (taxa retention). (**i**) Zeta_5_ of full community over age. (**j**) Best model fit to zeta diversity decline and taxa retention curves according to AIC. (**k**) Relationship between mean share of single community in % for a time-slices vs respective Zeta_5_ showing positive slopes with different slopes. (**l–m**) Zeta_5_ of communities and traits, p-values obtained from testing distribution differences to full community.

The 27, 25, and 23 ka BP time-slices exhibit notably higher zeta diversity values across all orders compared to later periods (Fig. 3a), indicating widespread taxa sharing among sites during LGM (28-20 ka). This trend is particularly evident in temporal patterns of Zeta₅ (taxa shared by at least five sites), which peaks during early LGM, drops during the early Deglacial, and shows another pronounced increase towards the mid-Holocene (Fig. 3i).

The shapes of the zeta decline curves differ between time-slices. Comparisons of models fitted to the zeta decline curves, evaluated using the Akaike Information Criterion (AIC), indicate that power law models generally provide a better fit than exponential models for the full assemblage, except at 14 ka BP where the difference between the exponential and the power law model is negligible (Fig. 3j and fig. S6). For zeta ratio (taxa retention) curves, strong asymptotic models generally provide the best fit for most time slices (fig. S7); however, modal models are optimal during the Bølling–Allerød period, and linear/segmented models prevail for the early Holocene (Fig. 3j).

Generally, normalized Zeta₅ values correlate positively with the mean percentage of the community in a time-slice, but the slope of the line varies between communities, being especially steep for forb tundra and comparatively shallow for shrub tundra (Fig. 3k). Normalized temporal Zeta₅ values of individual metacommunities vary distinctly compared to the full community. The forb tundra community forms a high proportion of the vegetation during LGM (Fig. 2a), which likely underlies the particularly high Zeta₅ values observed for this community, with about 30% of taxa consistently shared across at least five records (Fig. 3l). The shrub tundra community has elevated Zeta₅ values during early LGM and from approximately 13 ka onwards, while boreal forest Zeta₅ distinctly peaks in the early Holocene. Student’s t-tests revealed that the normalized Zeta₅ values for forb tundra, cushion plants, shrubs, and stress-tolerant taxa differ significantly (p < 0.1) from those of the overall community (Fig. 3m).

Zeta₅ values for individual sites range from about 15 to 95. Lake Levinson Lessing and Salmon Lake exhibit particularly high Zeta₅ values (Fig. 3c), suggesting notable consistency in taxa presence at these locations.

## Discussion

### Spatio-temporal dynamics of plant metacommunities in Beringia

Our analysis demonstrates that plant assemblages from all investigated sites across Beringia follow coherent temporal trajectories (Fig. 2c, **H01**, Fig. 4, and fig. S3) reflecting the cold and warm climate extremes during the LGM and the early Holocene in the Northern Hemisphere high-latitudes, respectively (*27*). We assume that spatially autocorrelated environmental changes enhanced both compositional synchrony and the magnitude of aggregated ecological responses across multiple sites

**Fig. 4.**
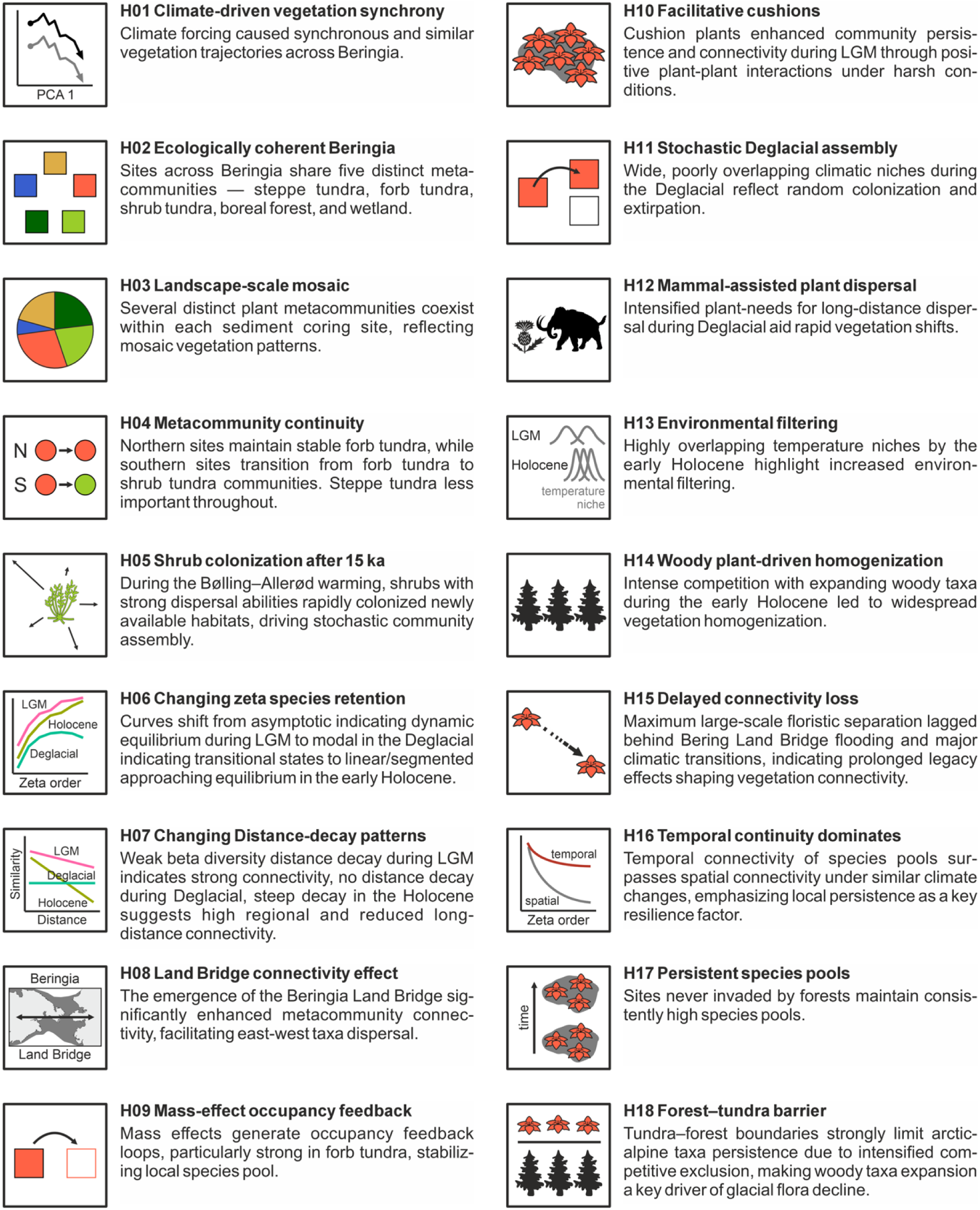
Emerging hypotheses distilled from our sedimentary ancient DNA analyses and exploratory data synthesis—intended for rigorous testing in subsequent studies. See discussion part for details.

(*28*). Changes in the community and trait composition (Fig. 2) further substantiate this interpretation, highlighting open habitats during the LGM, characterized predominantly by taxa with low-temperature niches, graminoid or cushion growth forms, ruderal CSR strategies, and limited dispersal capacities. The Bølling–Allerød warming (∼15 ka) marked a compositional shift towards taxa with warmer temperature niches, high dispersal capacity, increased shrub dominance, and the prevalence of stress-tolerant species, culminating in significant tree expansion toward the early and mid-Holocene (Fig. 2h). Accordingly, climatic traits explain a large proportion of the compositional variation, which is largely coherent with those explained by the functional traits (table S3). Collectively, these results highlight glacial-interglacial vegetation change as a substantial functional ecological reorganization (**H01**) at Beringia-scale, aligning with previous study results (*13*, *29*, *30*).

The observation that unconstrained ordination involving all sites and time-slices primarily reflects a temporal rather than spatial gradient suggests that ecological responses to climatic changes are embedded within an interconnected Beringian species pool (**H02**). Our network analyses of plant metabarcoding co-occurrence data identified five distinct communities (which we assume to represent plant metacommunities *sensu* Leibold et al. (*5*). We interpret each metacommunity as a set of interacting local communities interconnected across Beringia through dispersal, with each metacommunity occupying a distinct habitat. We assume that habitat patches of the different communities frequently co-occur at the landscape-scale within individual lake catchments (**H03**) and their sedaDNA traces are thus preserved in the lake sedimentary archives. The identification of metacommunities across multiple sites, rather than isolated within a single site, further supports viewing Beringia as an integrated ecological and environmental entity (**H02**). The generally high temporal and spatial zeta values support our inference of interconnected communities in space and time.

Paleoecological proxy-studies and biome niche modeling have highlighted a widespread, homogeneous mammoth steppe biome during the LGM (*10*, *31–33*). Similarly, our analyses identified a distinct **steppe tundra** community (Fig. 1) characterized by typical taxa known from modern Siberian steppe-tundra relics or central Asian steppes (*34*). However, in our records this steppe tundra metacommunity represents only a minor fraction of the glacial vegetation in Beringia. This apparent discrepancy might partly result from taphonomic limitations; dry upland habitats are likely underrepresented in sedimentary records, whereas moist wetland stands tend to be overrepresented. Furthermore, Poaceae may be systematically underrepresented in sedaDNA due to their less-developed root systems, with soil being the primary DNA source. Our strict 100% match criterion may miss extinct tundra-steppe taxa, but as shown by Courtin et al. (*20*), likely extinct species—comprising ∼5% over 30 ka—are generally rare and unlikely to have influenced our main quantitative patterns.

Our results indicate that the glacial vegetation in Beringia was a mosaic-like mixture of metacommunities (*34*, *35*). According to our reconstruction, most sites were dominated by **forb tundra** (**H04**), compositionally similar to the modern forb tundra prevalent in modern time-slice across northern sites (Fig. 1a). RDA_forb tundra_ yielded that, compared with other communities, climate niche is a less important trait but functional traits are more important (table S3).

The **shrub tundra community**, dominated by dwarf shrubs including many Ericaceae and *Salix*, closely reflects contemporary shrub tundra communities widely distributed across northern Beringia and circumpolar regions (*36*). Our findings reinforce earlier pollen-based studies documenting significant expansions of warmth-preferring shrubs (**H05**) during the Bølling–Allerød and early Holocene warming periods (*37–39*). Our data, in addition, indicate that climatic niches and dispersal capacities account for the largest proportions of variance in shrub tundra community composition, whereas other functional traits and specific site locations play comparatively minor roles (table S3). Nevertheless, regional distinctiveness remains relevant; for instance, *Boykinia richardsonii*, which is native exclusively to Alaska and the Canadian Arctic, underscores biogeographical uniqueness within these shrub tundra assemblages.

The **boreal forest** community comprising trees, erect shrubs, and aquatic plants appears distinctly underrepresented in sedaDNA from northern regions compared to pollen data, likely due to long-distance pollen dispersal and higher pollen productivity of woody taxa (*18*). The RDA_boreal forest_ reveals that growth form, plant life strategy (CRS), and site best explain the variance in its composition (table S3); this may indicate that the community composition was limited by variations in the specific environmental conditions between habitats rather than general climate.

The **wetland** community has a largely similar composition to the widespread modern Siberian lowland wetlands (*40*). Increases in *Sphagnum* and wetlands in general since the mid-Holocene align with moisture and circum-arctic peatland reconstructions (*41–43*).

### Changes in Beringian connectivity and causes

#### Enhanced ecological connectivity during the Last Glacial Maximum (LGM)

Our analyses of spatial diversity for single time-slices indicate significantly **enhanced ecological connectivity across Beringia** during the LGM. Elevated Zeta₅ highlights substantial species overlap across multiple sites, suggesting broadly distributed metacommunities *sensu* Leibold et al. (*5*), predominantly forb and steppe tundra in our study (Fig. 2a). The strong asymptotic form of the zeta retention curves (Fig. 3k) underscores a state of dynamic equilibrium (**H06**), typical for ecosystems characterized by evenly distributed species and balanced colonization-extinction dynamics, resulting in stable community compositions. Low beta diversity, along with minimal distance decay in community similarity (**H07**), supports extensive floristic connectivity and spatial homogenization at broad geographic scales (*44*).

We propose that the enhanced ecological connectivity observed during the LGM was at least partially facilitated by the establishment of the Bering Land Bridge (**H08**) (*7*). This interpretation is supported by our findings of initially high beta diversity between Ilirney18 (Chukotka, western Beringia) and site E5 (Alaska, eastern Beringia), which markedly declined after continuous terrestrial connectivity was established around 35 ka (Fig. 2i). This connectivity facilitated intercontinental migration and subsequent community homogenization. Additionally, the presence of the land bridge enhanced regional continentality, likely buffering against global temperature declines towards the LGM (*9*) and thereby promoting environmental stability and facilitating east-west ecological connectivity in Beringia.

Several biotic mechanisms likely contributed to the resilience of the vegetation communities, even under the harsh climatic conditions of the LGM. We suggest that **mass effects**—where immigration from productive habitats sustains populations in otherwise suboptimal environments (*4*)—played a particularly important role (**H09**). As anticipated, we observe a positive correlation between the relative proportion of each community and their corresponding Zeta₅ values. Steppe tundra and forb tundra communities, in particular, likely benefited substantially from the mass effect as indicated by the steep slope (Fig. 3j). We assume a positive feedback loop between productivity and community stability, enabling even tundra species with limited dispersal capabilities to persist at the landscape scale due to spatial proximity effects of nearby habitats. However, interpretations of the mass effect should be approached cautiously, as our analyses consider only relative community cover, not biomass, thus potentially overlooking open-land fractions.

Aside from the mass effect, **facilitative plant interactions** have been demonstrated to stabilize community composition (*45*) under severe environmental stress by buffering abiotic conditions and may act at the large spatial scale [in contrast to competition (*46*)]. Using the dataset analyzed here, Liu Y. et al. (*47*) highlight the occurrence of such interactions involving cushion-forming species like *Saxifraga oppositifolia*. This fits with our finding that cushion plants have a particularly high Zeta_5_ (significantly different from full assemblage) indicating strong facilitation in habitats with a high nurse plant proportion supported by a homogeneous composition (**H10**) (*48*), an effect which we not observe for the other non-woody groups like graminoids and other forbs. Furthermore, Liu S. et al. (*49*), employing sedaDNA-based ecological network analyses from our study sites, provide evidence for significant **mammal-plant trophic interactions** during this period. Mammalian herbivores notably modulated nutrient cycling processes, stabilizing vegetation structures by mitigating competitive exclusion and enhancing biodiversity (*50*).

#### Rapid turnover led to disconnectivity across all scales during the Deglacial period

The Deglacial period (19–11 ka BP) saw major ecological restructuring along with rapid warming (*51*). The early Deglacial (19–16 ka BP) is characterized by taxa with low temperature niches and low dispersal abilities (Figs. 2b and 2f), indicating a preference to invest in local survival under harsh conditions. This may explain the minimal Zeta₅ diversity values (**H06**), accompanied by high beta diversity which lacks the usual distance-decay pattern, i.e. even geographically close sites show large compositional differences (**H07**). Niche assessment finds the co-occurrence of taxa with different temperature preferences (Fig. 2b) indicating high stochasticity in the local community assembly (**H11**). Modal zeta retention curves between 14 and 18 ka BP (Fig. 3k) indicate high heterogeneity and rare-species dominance, reflecting ecological instability (*25*) and likely limited niche occupancy due to large distances from glacial refugia (**H06**). Collectively, these observations suggest reduced connectivity within metacommunities during this period.

Along with the Bølling–Allerød warming at 15 ka BP, taxa with strong dispersal abilities—particularly shrubs like *Salix*, *Betula*, and *Alnus alnobetula*—rapidly colonized newly available habitats (**H05**) [aligning with pollen records (*38*)], eroding the high beta diversity characteristic of the early Deglacial (**H07**) which is in accordance with metacommunity theory (*23*). Despite comprising over 60% of the vegetation, the shrub community’s Zeta₅ values remain relatively low and do not differ significantly from the overall community, indicating limited mass effects. At 13 ka BP, the exponential zeta decline curve suggests stochastic community assembly driven by frequent new colonization and taxa loss events (*25*).

The rapid expansion of shrub communities was likely facilitated by pre-existing tundra vegetation, where positive interactions and abundant nurse plants supported the establishment of incoming taxa (*47*). The strong plant–mammal dispersal links—particularly with mammoths—revealed by sedaDNA-based network analyses across Beringia (*49*) suggest that effective long-distance dispersal was a limiting factor in plant migration and therefore vegetation connectivity (**H12**) during the Deglacial. Moreover, rising sea levels during the Deglacial period gradually submerged the Bering Land Bridge, restricting intercontinental connectivity and reinforcing regional differentiation (*52*).

#### Holocene connectivity was low at the continental scale but high at the regional scale

The Holocene was characterized by the maximum expansion of boreal forests alongside widespread shrub communities, reflected by elevated zeta₅ values indicating strong connectivity between sites. The zeta diversity decline closely follows a power law model (substantially lower AIC compared to the exponential model; fig. S6), underscoring deterministic community assembly processes driven by niche- and trait-based factors (*25*). Particularly during the earliest Holocene period (*5*), narrow and highly overlapping temperature niches (Fig. 2d) strongly suggest niche sorting facilitated by intense environmental filtering (**H13**) (*5*). This intense niche filtering likely reflects low winter insolation, as shrubs and trees exposed above snow experience stronger winter constraints than herbs insulated beneath snow (*53*).

Subsequently, during the early to mid-Holocene (10–6 ka), the linear/segmented (i.e. linear with break points) shape of zeta retention curves (Fig. 3k and fig. S7) indicates progressive vegetation homogenization dominated by widespread taxa. This likely resulted from intensified plant–plant competition (*47*) and phylogenetic sorting (*21*) along with the woody taxa dominance (**H14**). Trophic mammal–plant interactions also intensified, now dominated by smaller mammals like Cervidae and Leporidae with limited foraging ranges and less disruptive effects than Pleistocene megaherbivores (*49*), contributing to reduced ecological heterogeneity at landscape scales. This synchronous expansion of mature, spatially coherent shrub communities was likely supported by well-developed seed banks linked to soil formation (*54*) and rodent-mediated seed dispersal (*49*).

The slope of the beta diversity distance-decay relationship notably steepens with the onset of the Holocene (Fig. 2m) indicating much higher connectivity at short rather than large distances. The flooding of the Bering Land Bridge around 11 ka BP (*52*) likely acted as a major driver of intercontinental disconnectivity. Additionally, intensified early-Holocene insolation likely enhanced north-south continentality, amplifying the broad-scale latitudinal temperature gradient and reinforcing regional floristic differentiation. However, the maximum large-scale floristic separation between eastern and western Beringia—indicated by increased beta diversity between Ilirney18 (Chukotka) and site E5 (Alaska)—as well as the steepest slope in the distance-decay relationship, occurred during the mid-Holocene, several thousand years after the opening of the Bering Strait (**H15**) and following the early-Holocene warming peak. Further research is required to confirm whether this delayed differentiation reflects a millennial-scale legacy effect, which is known to influence plant community assembly dynamics in this region (*55*).

### Local species pool stability - resilience vs sensitivity

Spatial differentiation clearly emerges along the second PCA axis of the full assemblage (fig. S3), reflecting an east-west geographic gradient across Beringia, consistent with constrained ordinations showing high site-driven compositional variance (Fig. 1d). We assume that the spatial differentiation only partly arises from regional postglacial climate differences, such as the more pronounced early-Holocene warming in eastern Beringia compared to western Beringia (*56–58*).

The primary reason for the observed patterns may relate to community resilience. Our analysis reveals that the median temporal Zeta₅ diversity calculated temporally within individual sites is 10%, approximately twice as high as the median Zeta₅ values calculated spatially across multiple sites within individual time slices. This notable difference is especially surprising given that the median temperature niche variation is quite similar (2.1°C temporally versus 2.3°C spatially; fig. S8). This observation suggests that temporal connectivity (continuity) is substantially stronger than spatial connectivity under similar climatic gradients (**H16**). Consequently, species appear to persist locally over time, showing lower broad-scale migration rates than expected based solely on their climatic niches. Such findings align with the concepts of extinction debt and delayed local extirpation, wherein taxa exhibit lagged responses to environmental changes (*59*). However, we acknowledge potential biases arising from confounding factors such as site-specific depositional environment stability, which typically varies less within sites than across spatially distant locations.

Sites that were never invaded by forests—specifically Salmon Lake, Lake Rauchagytgyn, and Lake Levinson Lessing—maintained a high species pool over tens of thousands of years, despite dominance by taxa with limited dispersal capacities (**H17**). This contrast is particularly evident in the divergent zeta decline curves observed between the tundra-dominated Lake Rauchuagytgyn and the neighboring forest-influenced Lake Ilirney (Fig. 3g). Despite their close geographic proximity, separated by only a single mountain range, these sites exhibit distinctly different floristic trajectories. The consistency of zeta decline curves between records from two separate Lake Ilirney basins indicates that these differences are not attributable to analytical uncertainty (Fig. 3g). Even following forest retreat during later Holocene stages, sites previously influenced by forests retained a smaller fraction of their historical site-specific species pools compared to earlier periods. We propose that at Lake Rauchuagytgyn, spatial stasis or landscape-scale spatial compensation (e.g., source-sink dynamics at scales equivalent to the lake catchment, meters to kilometers) significantly supported the resilience of the species pool (*28*). In contrast, at Lake Ilirney, insufficient local sources of tundra taxa during the early Holocene limited the ability to sustain a diverse forb–tundra community. Thus, fine-scale species persistence, together with migration processes, strongly influence the composition of the species pools. This pattern corroborates previous findings that the tundra–forest boundary represents a significant floristic barrier (**H18**), restricting the persistence of arctic-alpine taxa (*60*); it likely reflects sensitivity to interspecific plant– plant interactions, especially competitive exclusion (**H14**). Consequently, competitive exclusion associated with the expansion of woody taxa emerges as the primary driver of glacial flora decline (*47*), potentially contributing to global extinction events for certain taxa (*20*). The inability of invading boreal species to fully compensate for the loss of Arctic flora may result from disparities in their migratory capacities.

## Conclusions

Our study analyzed metabarcoding results from sedimentary ancient DNA across multiple sites in Beringia, delineating five distinct metacommunities—steppe tundra, forb tundra, shrub tundra, boreal forest, and wetland—by network analyses. The observed vegetation and trait dynamics reflected clear ecological responses to climatic variability: the LGM vegetation were mostly open steppe and forb tundra dominated by taxa with limited dispersal and ruderal traits; the Deglacial exhibited rapid turnover driven by expanding shrubs and woody taxa; and the Holocene featured stable forest and shrub tundra communities underpinned by deterministic assembly processes.

The metacommunity concept effectively contextualizes these observed dynamics. During the LGM, mass effects facilitated stable and homogeneous forb and steppe tundra communities by maintaining species populations across diverse yet interconnected habitats (*5*). The subsequent transition into the Deglacial period was marked by stochastic, distant dispersal-driven dynamics, resulting in community instability and high compositional turnover. The Holocene vegetation exhibited deterministic structuring, characterized first by clear niche differentiation and later on by stable intra-community interactions – intense competition – consistent with equilibrium metacommunity states (*28*). Thus, Beringian vegetation dynamics highlight the intricate ecological interplay between climatic fluctuations, dispersal mechanisms, and biotic interactions across glacial-interglacial cycles.

The identified long-term species pool stability, particularly evident at sites that were not affected by forest invasion, supports resilience despite the limited dispersal capabilities of dominant taxa. Sites such as Lake Rauchuagytgyn illustrate high local zeta diversity and low compositional turnover, contrasting distinctly with nearby Lake Ilirney, demonstrating how subtle landscape-scale processes influence temporal connectivity dynamics. Conversely, boreal forest sites like Bolshoe Toko and Billyakh exhibit lower diversity and connectivity, highlighting the sensitivity of these communities to changing conditions.

The implications for contemporary climate change are profound. Current observations of increased beta diversity suggest unstable community structures accompanying rapid environmental changes. In analog to the past, present woody expansion (*61*) significantly disrupts tundra diversity and stability. Conversely, ecosystems with a high prevalence of cushion plants demonstrate a stable and homogeneous composition, indicative of resilience under stressful conditions (*45*). Shrubs demonstrate rapid response capabilities and high connectivity, driven by effective dispersal mechanisms, facilitating their dominance across the landscape. In contrast, boreal forest communities, relying on southern taxa dispersal, have not yet established stable, fully mature ecosystems, highlighting their ongoing susceptibility under warming conditions.

This comprehensive assessment underscores the pivotal role of connectivity in shaping Beringian vegetation dynamics, emphasizing both spatial and temporal connectivity as key determinants of regional biodiversity. The metacommunity framework significantly advances our understanding of ecological assembly processes, highlighting the critical roles of connectivity, dispersal, and biotic interactions across extensive spatial and temporal scales, with direct implications for managing and predicting ecological responses to ongoing and future climatic changes.

## Materials and Methods

### Sites and datasets

Sediments from 20 lake cores across Siberia and the Alaska region were analyzed in this study. Among them, sedaDNA plant metabarcoding data from Salmon Lake, Hidden Lake, and Schrader Lake are newly reported in this study. References for the sediment records including age-depth models and sedaDNA plant metabarcoding are included in table S1.

### SedaDNA laboratory pipeline

Generally, all core splitting, sample subsampling, DNA extraction, purification, and amplification were performed following protocols described in Courtin et al (*20*). For published core datasets, modifications of the laboratory pipeline are described in the respective references (table S1). Newly reported cores (Salmon Lake, Hidden Lake, Schrader Lake) were analyzed as follows. Sediment samples were analyzed in the paleogenetic laboratories at the Alfred Wegener Institute, Helmholtz Centre for Polar and Marine Research, Potsdam, Germany. DNA was extracted from ∼3–5 g of sediment per sample using the DNeasy PowerSoil Max Kit (Qiagen), followed by a purification and concentration of DNA with GeneJet PCR purification Kit (Thermo Fisher Scientific). Each extraction batch contained 9 samples and one extraction control. DNA concentration of the samples was measured with Qubit Flourometer (Thermo Fisher Scientific) and was then adjusted to 3 ng/µl whereof 3 µL were added to each PCR reaction to amplify the P6 loop of the chloroplast *trn*L (UAA) intron using tagged g and h primers (*17*). PCRs were run in independent triplicates and a no-template-control (NTC) was added to each PCR batch. PCR products were then purified using the MinElute Kit (Qiagen) and pooled together. PCR-free library preparation and Illumina paired-end sequencing on a NovaSeq platform was performed by Genesupport Fasteris SA (Switzerland).

### Genetic data and bioinformatics

All 20 lake core datasets were newly assessed applying the following bioinformatic pipeline. After DNA sequencing, the raw sequencing files were checked, filtered, and taxonomically assigned using the OBITools software v3 (*62*). First, paired-end sequences were merged using *obi alignpairedend*, then sequences were demultiplexed according to the tag-sequences using *obi ngsfilter* and further dereplicated into unique sequences with *obi uniq*. PCR or sequencing errors were removed using *obi clean* and *obi ecotag* was used for the taxonomic assignment against the customized “SibAla_2023” database (*20*). Amplicon sequence variants (ASVs) which had a 100% identity to the “SibAla_2023” database were kept for the following analysis. The OBITools outputs were filtered to retain ASVs identified at least to the family level. ASVs assigned to the same species were merged, while ASVs assigned to the same taxon at higher taxonomic levels were distinguished by numeric suffixes appended to the taxon name.

Before PCR replicates for each sample were merged together, the replicability of the replicates was assessed via non-metric multidimensional scaling (NMDS) using the *metaMDS()* function from the vegan package in R (*63*). PCR replicates were excluded if they showed substantial divergence from other replicates of the same sample or failed to cluster with them, suggesting poor quality or inconsistent composition within replicates.

### Trait data LegacyPlantTraits 1.0

Growth forms were retrieved from open-access databases including BIEN (*64*) and TRY (*65*), as well as from publicly available sources such as Google Images, Wikipedia and eFloras. Species were classified into one of the following categories: bryophyte, fern, forb, graminoid, shrub, tree, submerged/floating aquatic.

Dispersal distance classes (ranging from 1 to 6) were primarily sourced from the database created by Lososová et al. (*66*). For species not covered in this dataset, we applied the classification scheme used by Lososová et al. (*66*) and Vittoz & Engler (*67*), with some adaptations to the original concept. Further details on the classification criteria and modifications have been deposited in Zenodo (*68*). The classification of species into dispersal distance classes was based on the dispersal mode, dispersal related diaspore features, plant height, growth form, and habitat.

Plant heights were collected from the BIEN (*64*), TRY (*69*), EOL (*70*), TTT (*71*), and BET (*72*) databases, as well as from online botanical resources including Wikipedia, eFloras, Flora of North America, the Global Biodiversity Information Facility (GBIF), and British Bryological Society. Plant height was calculated as the average of minimum and maximum values or as 65-75% of the maximum height.

Dispersal mode was divided into five classes: local non-specific dispersal, anemochory, zoochory, myrmecochory, and hydrochory. From the data compiled from the GIFT, TRY, LEDA Traitbase (*73*), and EuDiS (*74*) databases, modes were calculated. For species lacking data, dispersal syndromes were inferred based on the mode of genera and families, supplemented by targeted online searches. Dispersal related diaspore features were also manually collected from online sources (e.g. Wikipedia, eFloras) and the characteristics were used to validate and adjust dispersal mode data following the system used by Lososová et al. (*66*).

Habitat information was primarily derived from the BET database and further supported by online resources (e.g. British Bryological Society, Wikipedia, eFloras).

CSR plant strategies (*26*) were assigned using *StrateFy*, a globally calibrated calculator tool developed by Pierce et al. (*75*). This method classifies species into competitive (C), stress-tolerant (S), and ruderal (R) strategies based on three leaf traits: leaf area (LA), leaf dry matter content (LDMC = leaf dry weight / leaf fresh weight), and specific leaf area (SLA = leaf area / leaf dry weight). Trait data were gathered from the BIEN, GIFT, TTT, TRY, and LEDA databases, as well as from regional datasets including Díaz et al. (*76*), McIntosh-Buday et al. (*77*) (Central European flora), Jin Y. et al. (*78*) (Tibetan Plateau), E-Vojtkó et al. (*79*) (Pannonian flora), Waller et al. (*80*) (NE North American flora), and the China plant trait database version 2 (*81*, *82*). Median values of LA, LDMC, and SLA were used in the calculation of CSR percentages.

For species with no trait data present, CSR strategy was inferred from genus- or family-level averages. If no such estimates were available, mean values were used. For bryophytes, CSR strategies were roughly estimated using *r–K selection classes* from the BET dataset, following the conceptual links proposed by Barreto (*83*) and the approach of Grime and Pierce (*84*). In cases of ambiguous or missing data, the r–K class was used. Further details on assigning CSR classes are available in the supplementary materials (*68*).

All trait data processing was performed in the R environment v4.3.0 (*85*), using the following R packages: GIFT (*86*), BIEN (*64*), rtry (*87*), and tidyverse (*88*).

Functional trait data and detailed data collecting protocols are published on Zenodo as LegacyPlantTraits 1.0 (*68*).

### Temperature niches

We applied the “plantDNA_PDF/SDM” framework (*89*) to nine high-resolution cores from Lakes Billyakh, Bolshoe Toko, E5, Ilirney16, Ilirney18, Levinson Lessing, Rauchuagytgyn, Salmon, and Ulu, continuously covering the past 28 ka, to estimate overlapping climate niches of plant taxa from sedaDNA metabarcoding data. Modern plant occurrences derived from the Global Biodiversity Information Facility (*90*, *91*) were linked to contemporary climatic data (*92*) using MaxEnt species distribution models (*93*). The modeled species distributions were used to generate taxon-specific probability density functions (PDFs) along a site-specific summer temperature gradient of ±7.5°C around the observed modern conditions, using the CREST method (*94*, *95*). PDFs were set up at the taxonomic resolution of the genetic marker. Sample-wise sedaDNA metabarcoding plant spectra were used to generate abundance-weighted joint PDFs, and reconstructed optima and uncertainty ranges (50%) were derived from the *crest.reconstruct()* function output from the crestr R-package (*95*). The same pipeline was used to generate taxon-specific PDFs along an annual precipitation gradient of ±500 mm around the observed modern conditions in order to reconstruct precipitation (not shown).

To calculate site-specific temperature niches, time-slices without reconstructed summer temperatures were filled using the median reconstructed optima from the neighboring time-slices. Upper and lower reconstruction uncertainties were calculated as differences between the reconstructed optima and the upper and lower 50% uncertainty ranges. Summer temperature anomalies were calculated by subtracting the reconstructed optimum from time-slice 1 (2 ka) from all time-slices. For each time-slice the mean for all cores was calculated. A mean from the reconstruction uncertainties was calculated and subtracted from the reconstruction uncertainties from each time-slice. Average reconstruction uncertainties for each time-slice were calculated with a LOESS smoother with span = 0.25.

Temporal pair-wise temperature differences were calculated by comparing the reconstructed temperature anomalies for all time-slices at the location of each core with the *combn()* function in R (*96*). In addition, we calculated spatial pair-wise differences for each time-slice by comparing all cores within the respective time-slice. Moreover, we randomized the temporal and spatial pair-wise temperature differences by running 20 repetitions of five randomly selected temperature values.

### Statistical analyses

We kept only ASVs that occurred in at least 3 samples, in at least 2 cores, and with more than 100 reads resulting in a dataset with a total of 278,870,834 reads. As a data quality threshold, we kept only samples that included > 20 taxa.

The co-occurrence network analysis included all samples from all sediment records and applied the Louvain method, a multi-level modularity optimization algorithm that efficiently identifies community structures within large networks (*97*), using the *cluster_louvain()* function from the igraph R-package (*98*) with a resolution of 1.1. To facilitate visualization, taxa within each metacommunity were ordered by species contribution to beta diversity (SCBD), calculated from read abundance data using *beta.div()* function from the adespatial R-package (*99*) with the Hellinger distance metric (table S2).

We have divided the dataset into 2 ka time-slices and summed the reads for each taxon within each time-slice. We only kept time-slices that included more than 10 taxa which resulted in a dataset of 256 timeslices with in total 226,261,847 reads. Abundances of aquatic taxa (table S2) were square-rooted to account for their better chance of being recorded than terrestrial plants.

The samples of each sediment core dataset were then split into 2000-year time-slices. We employed Principal Component Analysis (PCA) to identify site-trajectories of compositional change for all 20 sediment cores. We only included taxa assigned to one of the five metacommunities in the PCA analysis and applied a Box-Cox chord transformation to the taxa reads. The PCA sample scores were filtered for the nine high-resolution cores and plotted core-wise. A LOESS smoother with span = 0.5 was applied as average for each time-slice (Fig. 2c).

All further analyses were run on the nine sediment records that cover the 2 ka time-slices back to 28 ka continuously. A Redundancy Analysis (RDA) was conducted to quantify the variance explained by supplied variables including information about site, climate niches, and morphological and functional trait abundance (% of taxa) on community composition using the *rda()* function from the vegan package (*63*). We converted the taxa reads into percentages using the *decostand()* function from the vegan package (*63*), applied a Box-Cox chord transformation (*100*) and filtered for taxa with a minimum abundance of 1% occurring in at least 20 samples to reduce the noise from rare taxa. RDA was conducted with 90 taxa and 124 samples for the full assemblage. We also ran an RDA on single taxa-rich boreal forests, forb tunda, and shrub tundra metacommunities. We ran a set of RDAs to extract the explained variance by all variable groups together (full model), solely and uniquely by each variable group. In addition, we ran RDAs with a set of forward selected variables from all variable groups using the *ordiR2step()* function from the vegan package (*63*). Results of the RDA assessing the unique contribution of the variable groups including the forward selected single variables are provided in table S3.

Alpha diversity, representing site species richness per 2 ka time-slice, was calculated for each site and time slice as the number of taxa. To ensure comparability, we applied rarefaction techniques from minimum reads of 20,614 using the *rarefy()* function from the vegan package (*63*). Rarefied richness showed similar results.

Beta diversity, quantifying compositional turnover between communities, was calculated using pairwise Bray–Curtis distances among Box-Cox chord transformed samples. We analyzed beta diversity patterns through distance-decay relationships, examining how community similarity decreased with increasing geographic distances. Analyses were performed using the *beta.pair.abund()* function from the betapart R-package (*101*). Geographic distances between the sites were calculated with the *earth.dist()* function from the fossil R-package (*102*). We fitted distance decay models to pair-wise assemblage similarity with the *decay.model()* function from the betapart package (*101*).

Zeta diversity was analyzed to quantify the retention of taxa across multiple sites within individual time-slices and across multiple time-slices at individual sites (*103*) using the *Zeta.decline.mc()* function from the zetadiv R-package (*104*). In addition, we conducted normalized zeta diversity analyses by dividing the zeta values for each sample by the total number of species for this specific sample. Following (*25*), we selected the “ALL” approach because it explicitly accommodates limited sampling at a continental scale, does not impose spatial or temporal constraints, and thus ensures robust comparability across space and time. The *Zeta.decline.mc()* function automatically returns Akaike Information Criterion (AIC) values for exponential and power law zeta decline curves, which we compared to determine the best-fitting model. Multiple candidate models were fitted to retention curves (linear, linear/segmented, modal, and asymptotic; fig. S7) and compared using the AIC, identifying the best-fitting model for each time-slice. Normalized zeta diversity decline with an order of 5 (Zeta_5_) was also calculated for the five metacommunities and the growth form, dispersal, and life strategy traits including the determination of the best-fitting models (exponential or power law) for the respective trait. Moreover, we ran Student’s t-tests with the metacommunities and traits subsets against the full assemblage dataset with the “less” alternative hypothesis (i.e. the full assemblage dataset has a smaller mean than the subsets).

All statistical analyses and visualizations were implemented using the statistical software R version 4.2.2 (*85*).

### Additional datasets

A forest cover raster file was derived from the ESA CCI Landcover time-series dataset v2.0.7 (*105*). It provides 22 globally gridded landcover classes, from which we extracted the classes 70 (needle-leaved evergreen tree cover) and 80 (needle-leaved deciduous tree cover) including their respective subclasses and merged them into a forest cover raster, which we used in Fig. 1a to illustrate modern forest distribution.

PMIP4 GLAC-1D ice-sheet data (*106–108*) was used to extract land masks to represent unflooded land masses in Fig. 1b which are flooded at present. The GLAC-1D bathymetry for floating ice was extracted from netCDF format using the *nc_open()* function from the ncdf4 R-package (*109*), filtered for values > 0 (i.e. representing land) and exported as regular 1k time-slice raster layers with the *rasterize()* function from the terra R-package (*110*) using the data’s original spatial resolution. Landmass rasters for the respective time-slices in Fig. 1b were selected, converted into polygons using the *as.polygons()* function from the terra R-package, projected into World Mercator projection and simplified with the *ms_simplify()* function from the rmapshaper R-package (*111*). For the 26-28 ka time-slice we used the landmask for 24 ka and for the 26-40 ka time-slice we used the “flooded landbridge” situation (*7*) at the modern (0 ka) time-slice.

## Acknowledgments

We used AI-assisted text editing (ChatGPT-4, OpenAI); all AI-generated text was reviewed and revised by the authors. We thank Cathy Jenks for assistance with English language editing. We thank Simon Fey for extractions and laboratory work carried out as part of an internship and Master’s thesis. We thank Janine Klimke, Sarah Olischläger, and Simon Fey for contributions to the palaeogenetic laboratory work.

## Funding

This study was funded by the European Research Council (ERC, Glacial Legacy, grant no. 772852 to UH), the German Research Foundation (DFG) Gottfried Wilhelm Leibniz Award (grant no. HE 2622/34-1 to UH), the Deutsche Forschungsgemeinschaft (DFG, German Research Foundation, grants no. 514539694 to UH), the United States National Science Foundation (U.S. NSF, grant no. 2303462 to DK), the German Federal Ministry for Education and Research (BMBF, grant no. 03G0859A to MM).

## Author contributions

U.H. conceived the study. K.R.S.-L., and U.H. supervised the lab work. U.H. designed and supervised the bioinformatic and statistical data analyses, that were implemented by T.B. L.F. collected the trait data supervised by U.H., Contributors provided knowledge about sediment core collection or knowledge about original metabarcoding datasets. U.H. wrote the initial draft of the manuscript with contributions from K.R.S.-L., Y. L., S.L., T. B. and L. Z. F.

## Competing interests

The authors declare that they have no competing interests.

## Data and code availability

Raw sequencing data is available under the European Nucleotide Archive (ENA) and the Dryad repository with links provided in the given citations in table S1. Bioinformatic data analysis scripts, respective metadata files and R scripts for the abundance data analyses are documented under: https://github.com/PolarTerrestrialEnvironmentalSystems/DynamicConnectivities_PlantMetacommunities (This github repository will be publicly available and converted into a Zenodo DOI after acceptance of the manuscript).

## Notes

### Competing Interest Statement

The authors have declared no competing interest.

### Summary of Updates

In this revised version, we corrected the reference formatting. The previous version contained Zotero-specific links that were only accessible in browsers with the Google Scholar PDF Reader plugin installed. To avoid confusion, we have now removed these internal links so that the references appear as plain text. No other changes to the manuscript content were made.

